# Improvement of human donor retinal ganglion cell survival through modulation of microglia

**DOI:** 10.1101/2024.07.06.602307

**Authors:** Volha V. Malechka, Monichan Hayes Phay, Josy Augustine, Emil Kriukov, John Dayron Rivera, Clemens Alt, Meredith Gregory-Ksander, Petr Baranov

**Author notes:** Contributed equally (Volha V. Malechka, Monichan Hayes Phay). **Financial Support:** NEI/NIH U24 (PB), GFF (PB), DoD FTTSA VRP (PB), Gilbert Family Foundation (PB).

## Abstract

Stem cell-derived retinal ganglion cell transplantation therapy offers a promising avenue for restoring vision in patients with significant retinal ganglion cell loss. The major challenge in this therapeutic approach is ensuring the survival of transplanted human donor retinal ganglion cells within the host retina. Here, we demonstrated the pivotal role of host retinal microglia and macrophages in the rejection, acceptance and survival of human donor cells. To identify the potential targets for treatment in microglia-retinal ganglion cell interaction, we assembled a single-cell atlas for mouse retina during development, aging, maturation and across neurodegenerative conditions. To our knowledge, this is the largest integrated atlas containing 1,053,629 cells including 36,080 myeloid cells. The downstream analysis highlighted phagocytosis among other known pathways involved in microglia activation, neuroinflammation and host retinal ganglion cell damage. We hypothesized that the same mechanisms are responsible for donor retinal ganglion cell removal. We showed that it is possible to improve the survival of human donor stem cell-derived retinal ganglion cells through modulation of the host retinal microglia. Pretreatment of human donor retinal ganglion cells with annexin V and/or soluble Fas ligand before transplantation led to a 2.5-fold increase in donor cell survival, including the outgrowth of axons targeting optic nerve head. Detailed analyses of the host mouse retina post-transplantation revealed morphological changes microglia and macrophages typical for activation in neuroinflammation. These findings indicate that soluble Fas ligand and annexin V treatment can be used to improve the success rate of transplantation of neurons within the retina and central nervous system.

## Introduction

The degeneration of retinal ganglion cells (RGCs) is a key factor in various eye disorders, including different forms of glaucoma, hereditary optic neuropathies, ischemic optic neuropathies, and demyelinating diseases, all of which result in progressive axonal degeneration and death of RGCs, ultimately leading to irreversible blindness. ^1–5^ Despite significant advances, the clinical needs for these retinal neurodegenerative conditions remain largely unmet. Consequently, a variety of therapeutic strategies are being actively explored, including cell replacement therapy, pharmacologic neuroprotection, and gene therapy. ^6–9^ Among these, stem cell-derived RGC transplantation therapy holds considerable promise for restoring vision in patients with RGC loss and axonal degeneration. ^10,11^ One of the primary challenges with this therapeutic approach is ensuring the survival of the transplanted human RGCs within the host retina. In the majority of neuron transplantation studies, the survival rate remains below 10% and is generally attributed to donor cell death due to shear stress and mechanical damage, lack of trophic factors, hypoxia, several cell death mechanisms and activation of innate and adaptive immune responses. ^12–14^ In this study, we explored the contribution of microglia, the major effector cell of the innate retinal immune system, to the survival rate of donor human retinal ganglion cells (hRGCs) following transplantation. ^15^

In the healthy and normally developed retina, microglia reside in the outer plexiform layer (OPL) and inner plexiform layer (IPL), where they perform essential functions in immune defense, homeostasis, and development. ^16^ When exposed to damaging stimuli, microglia become activated and migrate to the sites of retinal injury, where they can become neurotoxic and eliminate damaged and stressed RGCs. ^17,18^ We hypothesize that the similar process happens in host retina microglia and macrophages after transplantation of donor RGCs. To identify the potential targets for treatment in microglia-RGC interaction, we assembled a single-cell atlas for mouse retina during development, aging, maturation and across neurodegenerative conditions. To our knowledge this is the largest integrated atlas containing 1,053,629 cells including 36,080 myeloid cells. The downstream analysis confirmed previous observations of similar pathways in microglia activation and highlighted the role of phagocytosis as a universal response to damage. We hypothesize that during cell isolation and transplantation, donor hRGCs undergo significant stress, which leads to the externalization of pro-apoptotic signals on their cell surface. ^19^ These signals attract tissue microglia and macrophages to the site of injury and initiate phagocytosis of the transplanted cells. By blocking these signals and modulating the host retinal microenvironment, we aim to prevent microglia and macrophages from recognizing and eliminating the transplanted cells, thereby enhancing the survival of donor hRGCs post-transplantation.

Annexin V emerges as a potential candidate for modulating the retinal microenvironment and preventing microglia and macrophages from recognizing and removing donor hRGCs after transplantation. Annexin V is a calcium-dependent, membrane-bound protein with high affinity for phosphatidylserine (PS), an endogenous phospholipid and a component of the plasma membrane that becomes exposed on the cell surface during cellular stress and early apoptosis. Given the distressing nature of isolation and transplantation, donor hRGCs likely externalize PS under stress. Therefore, we hypothesized that pretreating donor hRGCs with annexin V to mask PS would prevent recognition and clearance by microglia and macrophages following transplantation.

Fas ligand (FasL), or CD95 ligand, is another possible candidate for modulating the retinal microenvironment and improving survival of transplanted donor hRGCs. FasL is a 40kDa type II transmembrane protein of the tumor necrosis factor (TNF) family that is capable of inducing apoptosis upon binding to the Fas receptor. ^20^ Within the eye, FasL can be expressed as a membrane-bound protein (mFasL), which is pro-inflammatory and pro-apoptotic molecule, or cleaved and released as a soluble protein (sFasL), which is non-inflammatory and non-apoptotic. ^21,22^ Previous studies demonstrated that FasL deficiency or intravitreal administration of sFasL protects RGCs from cell death in a mouse model of glaucoma. ^23^ Furthermore, sFasL has demonstrated therapeutic benefits in reducing inflammation in experimental models of keratitis^24^ and autoimmune disease. ^25^ Thus, we hypothesize that Fas-FasL interaction may protect donor hRGCs from immune attack by host microglia and macrophages, thereby improving graft survival in the host retina. Taken together, this study demonstrates that pretreating donor hRGCs with annexin V and sFasL before transplantation modulates the host microenvironment and improves graft survival in the host mouse retina. Additionally, our results provide the first quantitative analysis of microglia and macrophages morphology in the mouse retina following donor hRGC transplantation.

## Materials and methods

### Animals

All procedures conformed to the National Research Council’s Guide for the Care and Use of Laboratory Animals, the Association for Research in Vision and Ophthalmology (ARVO) guidelines and were approved by the Institutional Animal Care and Use Committee (IACUC) of the Schepens Eye Research Institute (SERI). All animals were purchased from Jackson Laboratory (Bar Harbor, ME). The breeding of animals was established in the Rodent Barrier Facility at the SERI. At the beginning of the experiment equal numbers of male and female *Brn3b^-/-^* and *Cx3cr1^GFP/+^* on a C57BL/6J background mice were group-housed in a temperature-controlled environment with 12-hour light/dark cycle and received a standard diet and water ad libitum.

### Organoid dissociation and isolation of donor hRGCs

Retinal ganglion cells (RGCs) were differentiated from H9:BRN3B:tdTomatoThy1.2-hESCs in 3D retinal organoid culture, as was previously described.^26–28^ Mature human retinal organoids were dissociated, and RGCs were isolated using magnetic microbead sorting for Thy1.2+ cells. Following cell isolation, RGCs were formulated in RGC media at 1 – 5 x 10^5^ cells/mL. Then, donor hRGCs were pretreated with 5 mg/mL annexin V (BioLegend, San Diego, CA), a calcium-dependent membrane-bound protein with high affinity to PS, and 0.2 mg/mL sFasL (R&D Systems, Minneapolis, MN), a non-apoptotic, non-inflammatory protein, for at least 20 minutes on ice. Following donor hRGC pretreatment, the cells were formulated for transplantation at 2 x 10^4^ cell/µL in RGC media.

### *In vivo* transplantation of donor hRGCs

The *Brn3b^-/-^* mice were anesthetized intraperitoneally with 100 mg/kg ketamine hydrochloride and 20 mg/kg xylazine hydrochloride solution diluted with saline. Pupils were dilated with 1% tropicamide ophthalmic solution (Bausch + Lomb, Bridgewater, NJ). Following tropicamide application, one drop of 0.5% proparacaine hydrochloride ophthalmic solution (Bausch + Lomb, Bridgewater, NJ) was applied onto the cornea for 5 minutes before transplantation. Up to 2 x 10^4^ donor cells per 1 µL were injected into one eye of *Brn3b^-/-^* mice using a beveled glass microneedle with 100 µm inner diameter at a 1 µL/min flow rate. Each mouse received a single subretinal injection of 1.0 µL of donor hRGCs pretreated with annexin V and sFasL during transplantation. Control group received a single subretinal injection of 1.0 µL of donor hRGCs with no annexin V or sFasL pretreatment. Following donor hRGC transplantation, one drop of GenTeal® Tears Lubricant eye gel (Alcon Laboratories, Fort Worth, TX) was applied onto the cornea to protect the eye from dryness. Mice were returned to the housing room and maintained on a standard 12-hour light/dark cycle for 3 days. On day 3 mice were euthanized, and eyes were immediately collected and fixed in 4 % paraformaldehyde (PFA) for 12 hours before immunohistochemistry analysis.

### Immunostaining of whole mount retinas

After fixation with 4% PFA was complete, retina samples were incubated in blocking buffer [10 % goat serum, 1 % bovine serum albumin (BSA), 0.1 % sodium citrate, 0.1 % Tween 20, and 0.1 % Triton-X in 1X phosphate-buffered saline (PBS)] containing goat anti-mouse antibodies in dilution 1:400 at 4°C for at least 12 hours overnight to limit nonspecific staining. Following blocking, retina flat mounts were incubated with Iba-1 primary antibodies (ab178846, Abcam, Boston, MA) in staining buffer (1 % BSA, 0.25 % Tween 20 and 0.25 % Triton-X in 1X PBS) for 72 hours at 4°C. Then, each retina sample was washed thrice for 15 minutes with washing buffer (0.1 % Tween-20, 0.1 % Triton-X in 1X PBS) at room temperature to remove any unbound primary antibodies before adding secondary antibodies. Secondary antibodies were prepared in dilution 1:500 in staining buffer, and the retina samples were incubated with secondary antibodies for 24 hours at 4°C. Then, each sample was washed with washing buffer thrice for 15 minutes at room temperature and then incubated with 1 μg/mL DAPI (4’,6-diamidino-2-phenylindole) in 1X PBS for 20 minutes at room temperature to stain the cell nuclei. Following final rinse with 1X PBS, retina samples were mounted and stored in the dark at room temperature before imaging. Images of retina mounts were acquired using confocal microscope (Olympus, Tokyo, Japan), and the Iba1^+^ microglia and macrophages were imaged within the OPL and IPL.

### Morphometric analysis of Iba1^+^ microglia and macrophages

The image of each retina flat mount was divided into four quadrants with one mid-peripheral region being a representative image of each quadrant; therefore, a total of four images per retina was used for quantitative analysis of retinal microglia and macrophages. The analysis was applied to quantify retinal microglia and macrophages and assess morphology of retinal microglia and macrophages according to metrics such as cell shape, ramification, and complexity. Morphology analysis of total number of Iba1^+^ retinal microglia and macrophages within the OPL and IPL was performed on skeletonized images of Iba1^+^ cells from the retinal flat mounts using previously published protocols. ^29^ Briefly, eight Z-stack images (two images per retinal quadrant) were captured at 20X magnification from one retina per animal and processed using ImageJ software (provided in the public domain). The images were binarized using the “adjust threshold” function and skeletonized using the “skeletonize” function. The plugin “Analyze Skeleton” was then employed to generate the number of branches, number of junctions, and average branch length of the microglia and macrophages in the retina flat mounts. All morphometric parameter values were divided by the total number of cells per unit area of retinal tissue in mm^2^.

### Single-cell RNA-sequencing data analysis

Data analysis was performed using R v.4.3.1 in RStudio 2023.06.1 Build 524. To perform reference mapping and multi conditional RGC/myeloid studies, we compiled multiple publicly available datasets for the conditions such as mouse healthy developing retina, mouse healthy adult retina, mouse optic nerve crush model, and mouse glaucoma model retina. The complete list of publicly available datasets used in this study is available in **Supplementary Table 1**. Cell ranger outputs that include matrix, features, and barcodes were obtained from public GEO repositories, and further processed using Seurat v. 4.3.0.1. (Hao Y et al. Nat Biotechnol, 2023; Hao Y et al. Cell, 2021) Quality control metrics included filtering by nCount_RNA, nFeature_RNA, percent.mt, percent.rb. Doublets were removed using the DoubletFinder algorithm. (McGinnis CS et al, Cell Syst, 2019) Each dataset was processed independently for the quality control and data processing, followed by conditional integration that was performed using multiple approaches: 1) default Seurat integration, 2) Seurat RPCA-based integrated, 3) Harmony, (Korsunsky I et al. Nat Methods, 2019) 4) scVI. (Lopez R et al. Nat Methods, 2018) The selection of method was based on the number of batches, batches variation, and the number of cells. Upon the integration, we subset RGC/myeloid from each condition and reintegrate the resulting subsets into multi conditional RGC and myeloid atlases. Further downstream analysis was performed using Seurat and escape v.1.10.0 packages. Pathways for GSEA pathway analysis were obtained from Molecular Signatures Database.

## Results

### RGC survival is improved following blocking phosphatidylserine residues with annexin V on donor hRGCs

Annexin V pretreatment led to the increase in transplant success: 92% (11 out of 12 recipients) had >1% (200) donor cells at day 4, compared to only 55% (6 out of 11) in the control group. We also observed 2.5-fold change in number of donor RGCs per eye: with mean numbers at 1,280 cells in pretreated vs. 520 cells in control. In both groups grafted donor RGCs displayed extensive neurite outgrowth (>150 mm) and axon projections (>1,000 mm) (**Figure 1A, B**). Furthermore, in the annexin V pretreated group, four out of twelve transplants exhibited neurite outgrowth into the optic nerve head, compared to only one out of eleven transplants in the control group (**Figure 1C**).

**Figure 1.**
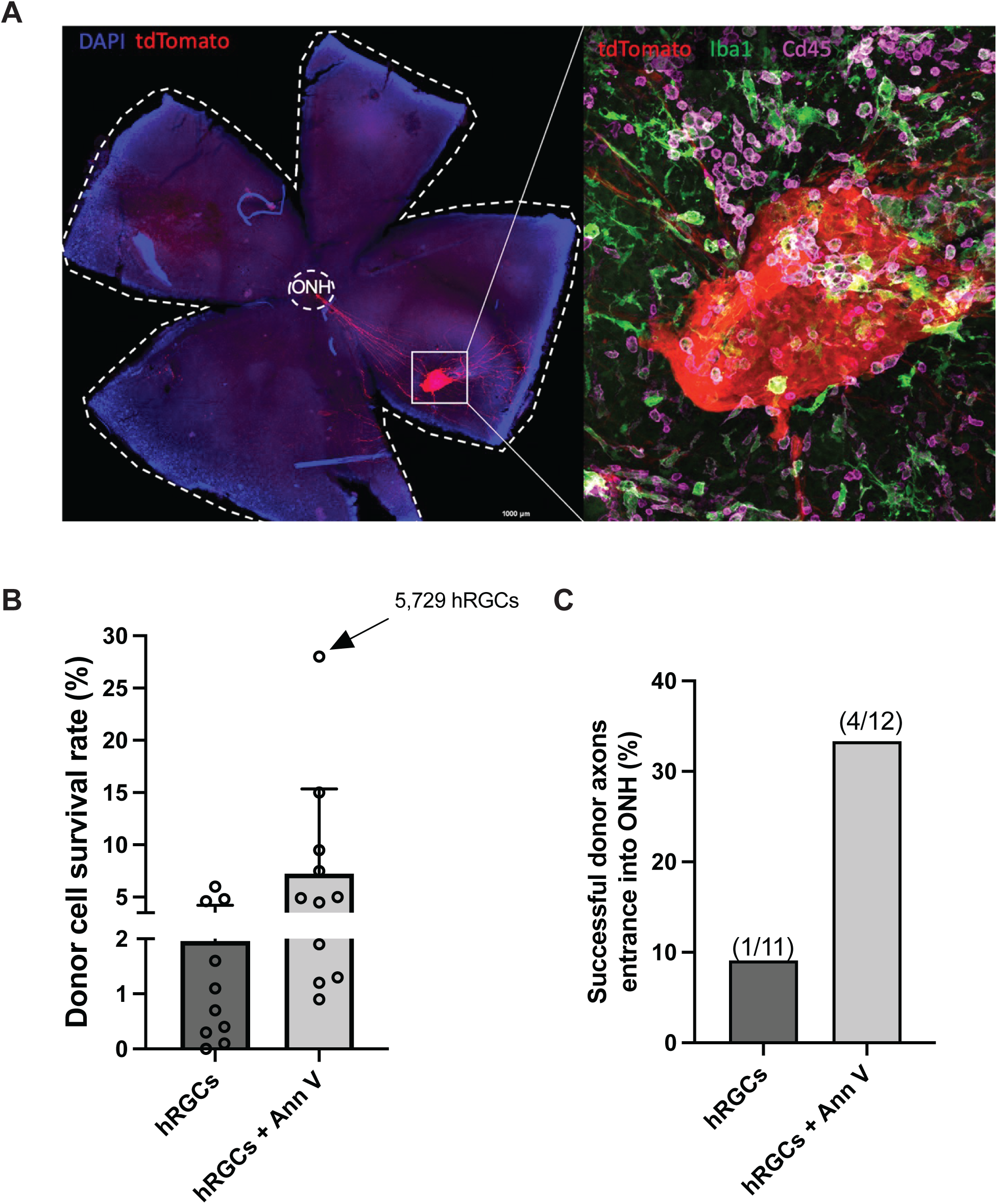
Donor cell survival is improved by blocking phosphatidylcholine (PS) residues on human donor retinal ganglion cells (hRGCs) with annexin V. (A) Donor hRGCs survive and send their projections to the optic nerve head (ONH). (B) Donor hRGCs survive better after pretreatment with annexin V. (C) Donor hRGCs enter ONH more successfully after pretreatment with annexin V.

### Host microglia and macrophages are activated after donor hRGC transplantation in *Cx3cr1-GFP* mice

To study microglia activation *in vivo*, we imaged the retina repeatedly in *Cx3cr^GFP/+^* mice by scanning laser ophthalmoscopy (SLO) ^30^ (**Figure 2**). On day 1 post-transplantation, host retinal microglia and macrophages become activated and initiate displacement toward injection site to remove the transplanted donor hRGCs (**Figure 2A, 2B**). By days 2 and 3, microglia and macrophages begin eliminating donor hRGCs, occupying the area of damage, and by day 7 all donor hRGCs are removed and processed by microglia and macrophages (**Figure 2A, 2B**).

**Figure 2.**
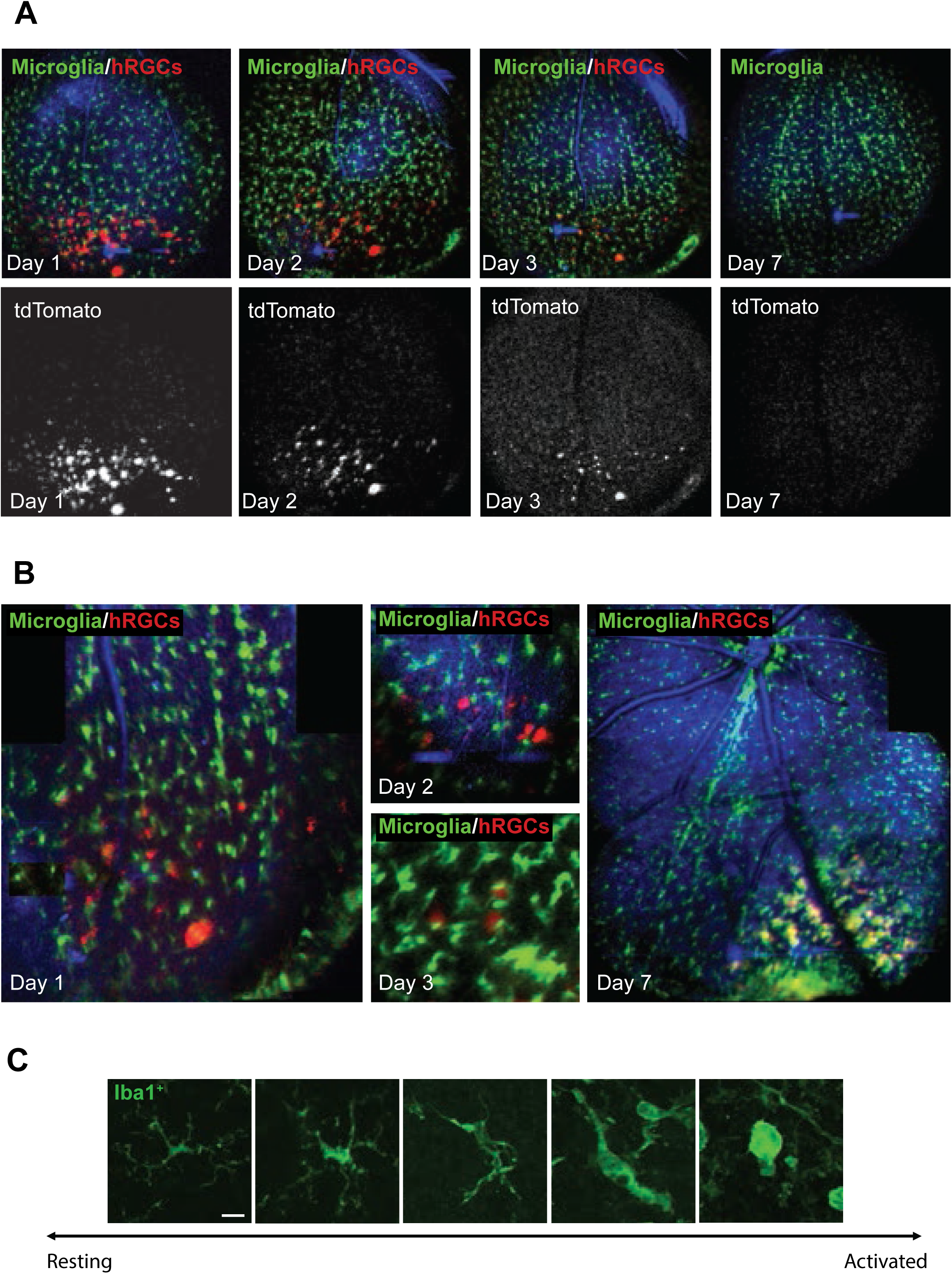
Studying microglia activation *in vivo* in the retina of *Cx3cr1-GFP* mice after donor human retinal ganglion cells (hRGCs) transplantation. (**A**) Imaging *in vivo* by scanning laser ophthalmoscopy (SLO). Image on day 1 represents injected donor tdTomato^+^ hRGCs (red) and host microglia cells (green) after transplantation. Note that by day 7 all injected donor tdTomato^+^ hRGCs are eliminated by activated host microglia cells. (**B**) Interaction between host microglia and donor hRGCs on a single cell level. (**C**) Morphological characteristics of a microglia cell. Scale bar: 5 μm

Retinal microglia and macrophages rapidly respond to physiological changes and demonstrate distinct morphology based on their state of reactivity. As shown in **Figure 2C**, activated microglia and macrophages retract their processes and round their cell bodies, whereas in their homeostatic state they exhibit small cell bodies with ramified processes.

### Donor hRGCs are eliminated by primary microglia *in vitro*

To determine if donor hRGCs are eliminated by microglia *in vitro*, we co-cultured primary microglia with human donor RGCs (**Supplementary Figure 1A**). We have observed that primary microglia become activated on day 1, move towards donor hRGCs, and begin eliminating them by day 1 (**Supplementary Figure 1B**). This finding is consistent with our observation *in vivo*, demonstrating that donor hRGCs are recognized and eliminated by microglia in the host microenvironment. Therefore, modulating this microenvironment might help suppress microglia activity and improve the survival of donor hRGCs.

### Activation of host microglia and macrophages is modified following exogenous pretreatment of donor hRGCs with annexin V and sFasL

To investigate whether the host retinal microenvironment, where microglia and macrophages play an essential role in active surveillance can be modulated, we pretreated donor hRGCs before transplantation. Following dissociation and isolation of donor hRGCs, we pretreated the cells with 5 mg/mL annexin V and 0.2 mg/mL sFasL for at least 20 minutes on ice. Following transplantation, our results indicate improved survival and axonal elongation in donor hRGCs pretreated with annexin V and sFasL as compared to non-pretreated cells (**Figure 3A, 3B**).

**Figure 3.**
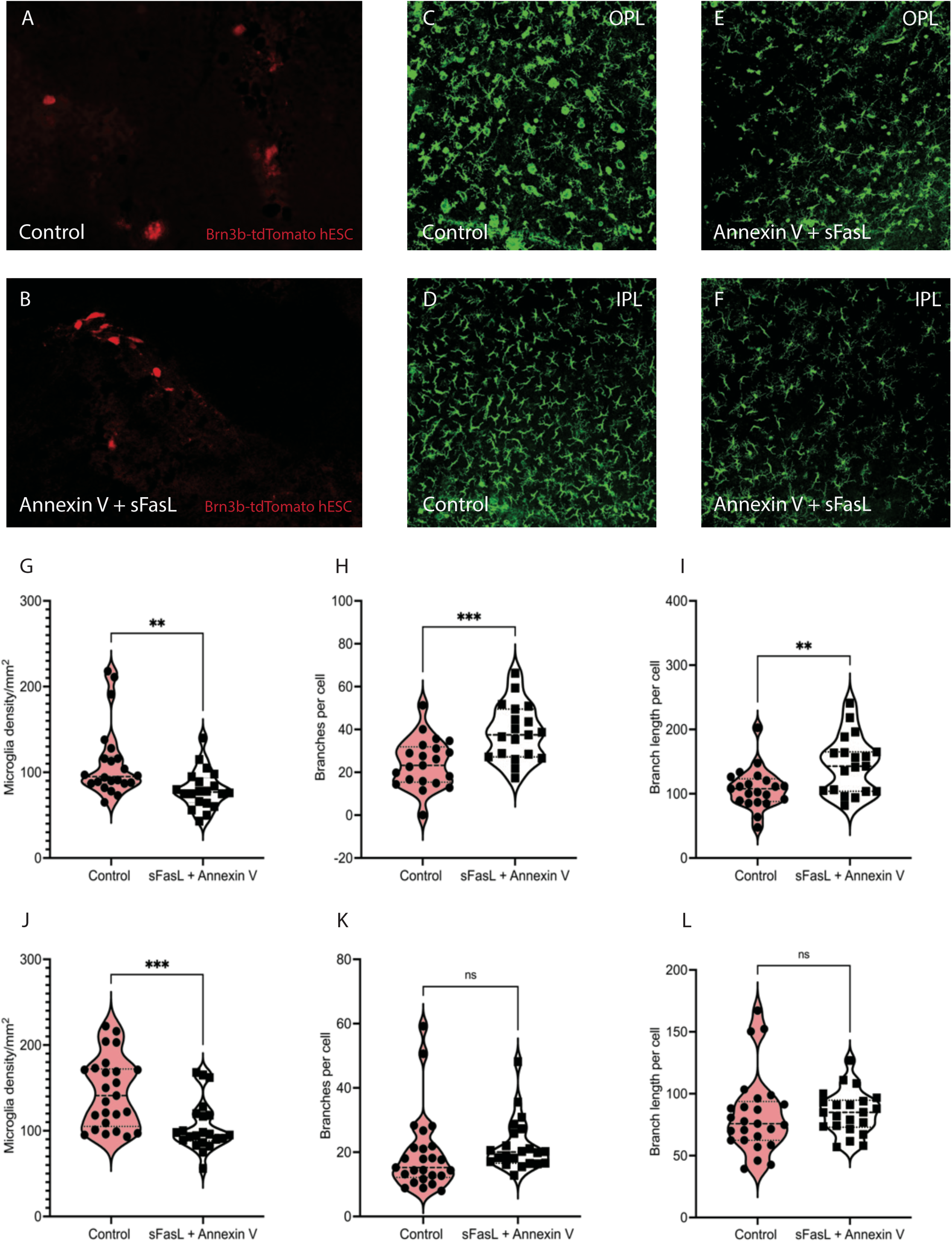
Pretreatment of donor hRGCs with annexin V and sFasL inhibits microglia activation and increases survival of donor hRGCs after transplantation. (**A, B**) Donor hRGCs survive better after pretreatment with annexin V and sFasL. Activation of Iba1-positive microglia and macrophages in the OPL (**C, E**) and IPL (**D, F**) of the retina in the control group and group pretreated with annexin V and sFasL before donor hRGC transplantation. Microglia density (**G, J**), number of branches per microglia and macrophage cell (**H, K**), branch length per microglia and macrophage cell (**I, L**) in the OPL (**G, H, I**) and IPL (**J, K, L**) of the retina in the control group and group pretreated with annexin V and sFasL. hRGCs, human retinal ganglion cells; IPL, inner plexiform layer; OPL, outer plexiform layer; sFasL, soluble Fas ligand.

Microglia are resident cells in the retina localized in two distinct layers within OPL and IPL in the intact retina. ^31^ To explore the effect of pretreating donor hRGCs with annexin V and sFasL on microglia and macrophages, we analyzed Iba1^+^ cells within OPL and IPL after donor hRGC transplantation. At 3 days post transplantation of donor hRGCs pretreated with Annexin V and sFasL, microglia and macrophages displayed homeostatic morphology with prominent and longer processes as compared to the control group with no pretreatment, where microglia and macrophages displayed a more reactive phenotype with rounded cell bodies and retracted processes (**Figures 3C-F**). Interestingly, we also found that microglia and macrophages become more activated in the OPL as compared to the IPL after subretinal delivery of donor hRGCs (**Figures 3C-F**). Moreover, the total number of microglia and macrophages was reduced in the pretreatment group as compared to the control group with no pretreatment (**Figures 3G, 3J**), again with more significant changes observed in IPL. Additionally, the number of branches per microglia and macrophage was significantly (*p*<0.01) increased in the pretreatment group, indicating less activation when compared to the control group (no pre-treatment) (**Figures 3H, 3K**).

Moreover, microglia and macrophages in the OPL exhibit a significant difference in branch number per cell between pretreated and control groups (**Figure 3H**), while those in the IPL showed lower branch numbers per cell (**Figure 3K**). Similar trend is observed in the branch length per microglia and macrophage cell with longer and more ramified branches in OPL that in IPL, suggesting that microglia and macrophages are less activated in the OPL than IPL after donor hRGC transplantation (**Figures 3I, 3L**). In summary, pretreating donor hRGCs with annexin V and sFasL leads to morphologic changes in microglia and macrophages within the OPL and IPL of the mouse retina.

### Characteristics of RGCs and microglia vary in the conditions of healthy developing, adult, acute and chronic post-injury retina

We built an atlas of microglia and macrophages from healthy developing retina, adult retina, optic nerve crush (ONC) retina, and microbeads-induced glaucoma model retina (**Figure 4A**) to better understand the process of neuroinflammation and the interaction of activated microglia with the retinal neurons. While some condition-specific states are present for developing microglia/macrophage, most clusters overlap between the conditions. However, the differences are observed when assessing the classical microglia-related genes expression (**Figure 4B**), where microglia and macrophages in the developing retina are mostly characterized by strong proliferation, expression of *Igfbpl1*, (Pan L et al. Cell Reports, 2023) *Ager*, and *Igf1*, and healthy adult retina state is profiled by *Stab2* and *P2ry12*. In ONC retina, we first observe low expression of *P2ry12*, a classic homeostatic microglia marker, that tends to increase over time. In addition, early timepoints in ONC retina are characterized by *Stab1* and *Ccl4* with higher expression of *Ccl5* week 2 post-ONC. Interestingly, we observe *Cdc20* and *Mki67* expression on days 2 and 4 post-ONC and an increase in *Apoe* expression on day 4 post-ONC. Retina in microbeads-induced glaucoma model is characterized by *Timd4* and *Olr1* expression demonstrating the pattern of increased *Cd74* expression compared to developing and healthy adult retinas. FasL expression is the lowest in developing retina, whereas the highest expression of FasL is observed in healthy adult and 2 weeks post-ONC retinas. These patterns demonstrate a complicated scheme of microglia activation between different conditions. To further inspect the conditional changes, we performed pathway analysis using the pathways related to microglia activation (**Figure 4C**). As expected, we observe myeloid progenitor cell differentiation to be the highest in developing retina. Within all pathways related to microglia activation, we see healthy adult condition to have the lowest enrichment score when compared to ONC and microbeads-induced glaucoma model conditions. The number of differences exists between microglia activation in ONC and microbeads-induced glaucoma model. ONC retina is characterized by higher microglia proliferation, migration, and differentiation, whereas microbeads-induced glaucoma model condition demonstrates a higher enrichment score in acute inflammatory response and innate immune response activation.

**Figure 4.**
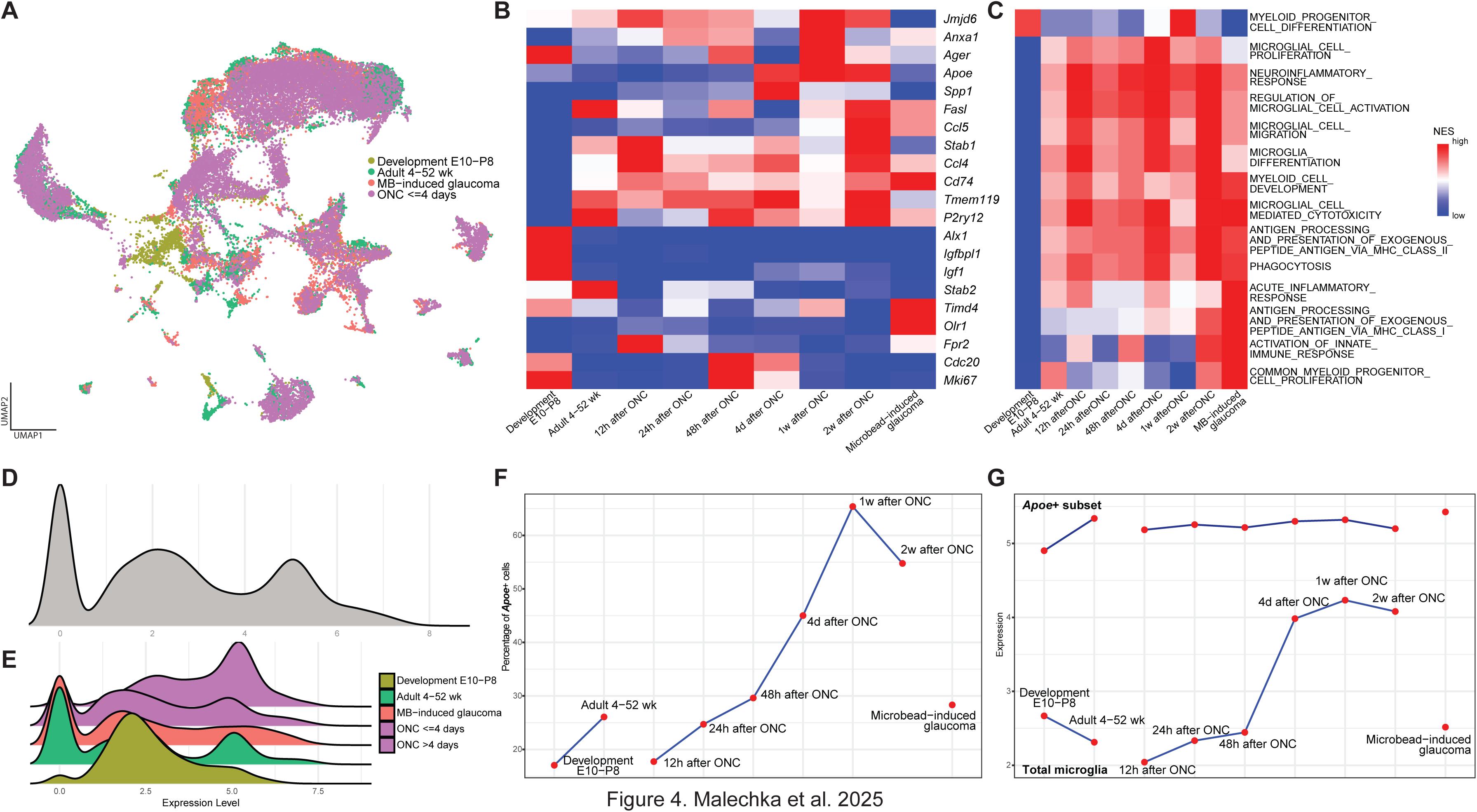
(**A**) Integrated dataset of mouse developing (E10-P8), healthy adult (4 – 52 weeks), microbeads-induced glaucoma, and optic nerve crush retinal myeloid population. (**B**) Heatmap demonstrating expression patterns of classical microglia-related genes between the conditions. (**C**) Pathway analysis demonstrates condition-specific, and microglia activation related signaling patterns between the conditions. Distribution of *Apoe* expression in the integrated dataset (**D**) and between the conditions (**E**). (**F**) Chart highlighting the differences in percentage of *Apoe^+^* (expression >= 4) cells between the conditions. (**G**) Dynamics of Apoe expression between the conditions for the total dataset (bottom) and *Apoe^+^* (expression >= 4) cells. E, embryonic day; P, postnatal day.

As we observe such drastic differences in microglia activation between the conditions, we explored the role of *Apoe,* one of the most variable markers in the conditions studied, in microglia activation. Overall, we observed three peaks in *Apoe* expression in the integrated dataset. The first peak shows *Apoe* cells, while the other two highlight *Apoe^+^* cells that for simplicity we call *Apoe^dim^* and *Apoe^bright^* (**Figure 4D**). Interestingly, the distribution of *Apoe* expression is different between the conditions. Microglia in developing retina demonstrates almost complete absence of the first peak of *Apoe^-^* cells, and microglia in ONC retina >4 days has the distribution shifted towards higher expression compared to ONC <=4 days condition (**Figure 4E**).

We further subset the *Apoe^bright^* population with the threshold of expression set to ≥ 4. We observe multiple trends in change of *Apoe^bright^* population: 1) the population is growing from healthy developing to healthy adult condition; 2) in the ONC timeline, the population is growing up to week 1 post-ONC, then decreasing; 3) glaucoma model condition population is similar to healthy adult retina (**Figure 4F**).

To understand the role of *Apoe* in microglia activation and to establish, whether it is a marker of increased microglia activation or a marker of cells undergoing transition from homeostatic to activated state, we compared datasets of average *Apoe* expression within two populations - total integrated microglia/macrophage population and *Apoe^bright^* population (**Figure 4G**). Noticeably, *Apoe* expression does not change in *Apoe^bright^* population. For the total integrated microglia/macrophage population, average *Apoe* expression decreases from developing to adult condition and is similar between the adult and glaucoma model conditions. In the ONC condition, it increases up to week 1 post-ONC.

In the same manner we integrated the multi conditional (developing, adult, and ONC) data for RGCs (**Figure 5A**). We analyzed canonical “eat-me”, “do-not-eat-me”, and “find-me” signaling markers between the conditions (**Figure 5B**). Healthy developing RGCs can be profiled by *Lrp8* and *Rab5a* expression. Healthy adult RGCs express *Rab5a*, however, they lose *Lrp8* expression. They are also characterized by *Cd93*, *Timd4*, *Stab2*, and *Tulp1* expression, demonstrating a similar pattern to 2 weeks post-ONC condition. The timeline of post-ONC condition can be separated into two functional timepoints, such as <=4 days and >4 days, where <=4 days RGCs are positive for *Rab7*, *Cd47*, *Adgrb1*, and *Cx3cl1*, while >4 days are positive for *Rab35*, *Gas6*, *Plscr3*, and *Spp1*. Noticeably, RGCs stand separately and are positive for the majority of “eat-me” markers week 2 post-ONC.

**Figure 5.**
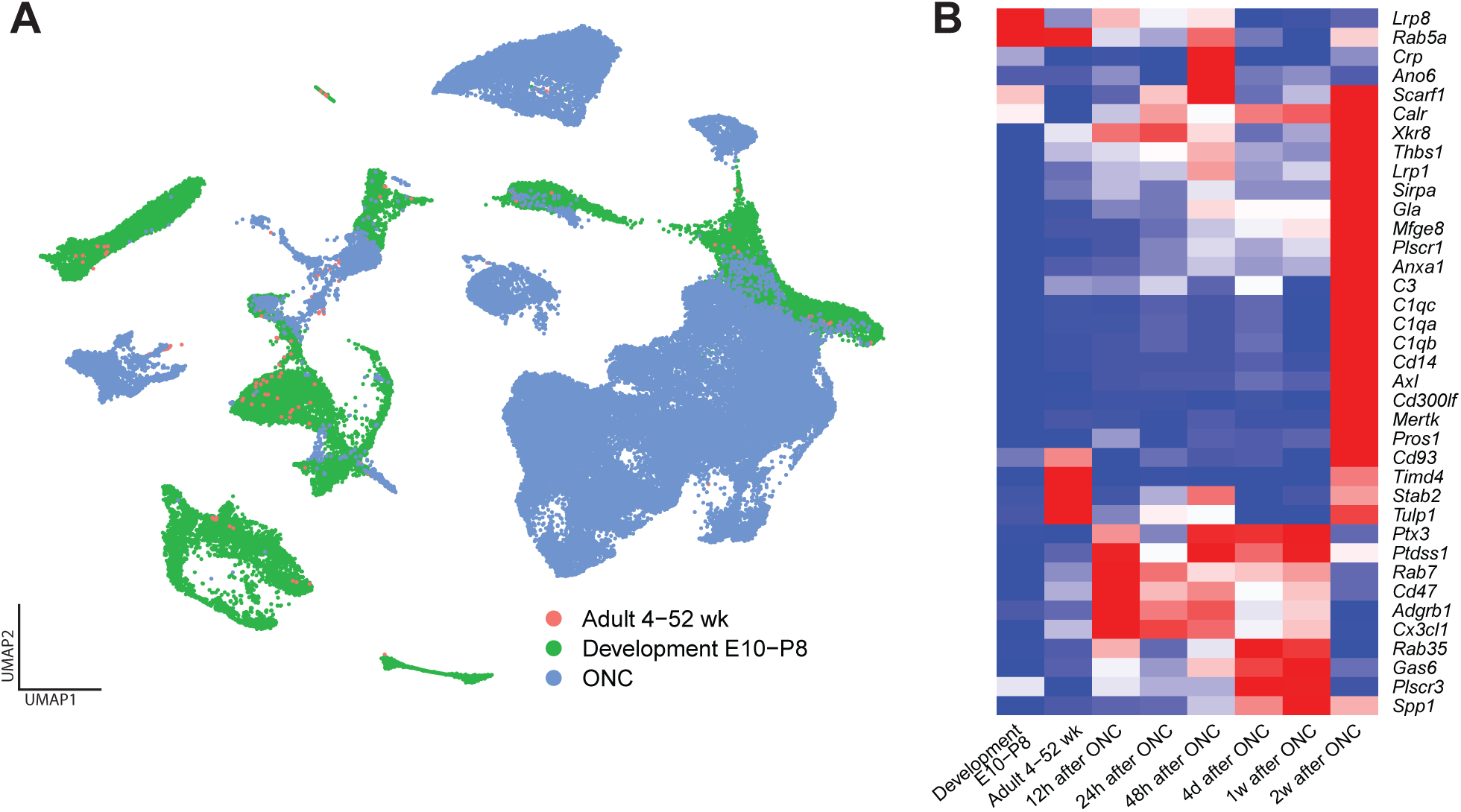
(**A**) Integrated dataset of mouse developing (E10-P8), healthy adult (4 – 52 weeks), and optic nerve crush retinal ganglion cell population. (**B**) Heatmap demonstrating expression patterns of known “eat-me”, “do-not-eat-me”, and “find-me” genes between the conditions. E, embryonic day; P, postnatal day.

## Discussion

Optic neuropathies and retinal neurodegenerative diseases, such as glaucoma, diabetic retinopathy, and age-related macular degeneration, result in progressive damage of retinal ganglion cells (RGCs). ^32^ If left untreated, these diseases lead to irreversible blindness. For the patients with optic neuropathies and neurodegenerative diseases of the retina, cell replacement therapy ^33–35^ and treatment with neuroprotective agents are promising strategies to halt disease progression and preserve vision.^36,37^ One of the key steps for improving cell transplantation success in the retina is to define the factors that positively affect survival of donor RGCs and to modulate the host microenvironment ^38^ by enhancing the beneficial effect of these factors on donor cells.

The retina is populated by specialized immune resident macrophages called microglia. Normally, retinal microglia reside within the outer plexiform layer (OPL), the inner plexiform layer (IPL), and the retinal ganglion cell (RGC) layer. Microglia represent one of the main barriers the donor cells need to overcome for successful survival and integration within the host microenvironment, as microglia are responsible for elimination of antigens that threaten the retinal integrity. ^38^

Microglial activation in the retina is a common feature in various retinal neurodegenerative conditions, including glaucoma and other optic neuropathies. Multiple studies ^39^ demonstrated the similarity of the activation process across mouse models. To better understand the microglia-RGC interaction and to identify potential therapeutic targets we have built an integrated atlas of mouse retina across conditions. These include developing, healthy adult, optic nerve crush, and microbead-induced glaucoma model. To our knowledge this is the largest integrated atlas containing 1,053,629 cells including 36,080 myeloid cells. The downstream analysis highlighted the similarities of the activation process, including the role of the phagocytosis in the elimination of the retinal neurons.

Prior studies demonstrated that phosphatidylserine (PS), a phospholipid that is localized to the inner part of the cell membrane, is crucial for identification of apoptotic cells by macrophages. ^40,41^ Considering donor RGCs experience a substantial stress during dissociation, isolation, and transplantation events, it is predicted they might generate pro-apoptotic signals and externalize a PS signal on the cell surface, allowing them to be recognized and cleared by retinal microglia and macrophages. ^42^ Annexin V, a calcium-dependent membrane-binding protein, efficiently and specifically attaches to PS, thus, representing a selective probe for recognition and quantification of apoptotic cells. ^43–45^ By pretreating donor RGCs with annexin V just after isolation and right before transplantation events, a PS signal on the cell surface of donor RGCs can be blocked, hence preventing donor RGCs from being recognized and attacked by retinal microglia and macrophages.

In addition to blocking the PS signal with annexin V, we previously demonstrated that treatment with sFasL is also neuroprotective and prevents death of RGCs in chronic and inducible mouse models of glaucoma. ^23,46^ Herein, we determined the combination of annexin V and sFasL significantly improves survival of donor hRGCs in the host mouse retina. FasL is a type II transmembrane protein that exists as both a membrane bound protein (mFasL) that is pro-inflammatory and pro-apoptotic, and a soluble form (sFasL) that is cleaved from the cell surface and released as a soluble fragment that is non-inflammatory and non-apoptotic and can antagonize the activity of mFasL. ^22,23,47^ In the healthy eye, the anti-apoptotic and anti-inflammatory sFasL is the dominant isoform expressed, ^46^ but with disease there is a shift and the pro-inflammatory and pro-apoptotic mFasL is the dominant isoform expressed in a variety of ocular pathologies. ^46,48–50^

In inducible and chronic models of glaucoma it has been shown that mFasL mediates destruction of RGCs either by directly killing RGCs and/or by inducing the production of pro-inflammatory chemokines or other mediators in Fas^+^ glial cells and thereby causing neurotoxic inflammation. ^23,46,51^ By contrast, intravitreal AAV2-mediated gene delivery of sFasL provided complete and long-term protection of RGCs and their axons. ^23,46^ Importantly, the sFasL-mediated neuroprotection of RGCs also correlates with reduced microglia activation and reduced production of pro-inflammatory mediators. Therefore, sFasL could be improving survival of donor hRGCs by blocking FasL-mediated apoptosis of RGCs and/or by blocking FasL-mediated activation of glia and the induction of neurotoxic inflammation.

Normally, in the healthy eye, retinal microglia display a highly ramified morphology with a small cell body and multiple branching processes. When responding to inflammation or injury, microglia retract their processes and present round cellular morphology. Although microglia are the most abundant immune cells in the retina, the retina also contains a population of perivascular macrophages that are located within the glia limitans of retinal vasculature. ^52^ The glia limitans acts as a physical barrier against unwanted cells that attempt to enter the retina. During the subretinal delivery of donor hRGCs, the microinjury to small retinal blood vessels may occur causing the leakage of perivascular macrophages to the site of injection. Both microglia and macrophages express the ionized calcium-binding adaptor (Iba1) molecule, a 17-kDa actin-binding cross-linking protein that is upregulated during activation of these cells. ^53^

Here, we investigated the morphology of Iba1^+^ microglia and macrophages following pretreatment of donor hRGCs with annexin V and sFasL before transplantation. We found that transplantation of donor hRGCs leads to local activation of host retinal microglia and macrophages causing them to move towards the injection site to eliminate donor hRGCs. Pretreatment with annexin V and sFasL resulted in reduced activation of microglia and macrophages, with Iba1^+^ microglia and macrophages displaying a more ramified morphology with longer processes (**Figures 3E, 3F**). In contrast, the Iba1^+^ microglia and macrophages in the control group exhibited a round ameboid morphology with retracted processes (**Figures 3C, 3D**).

Our results demonstrate that modulating the host retinal microenvironment through the microglia manipulation can improve the survival of donor hRGCs after transplantation. We believe it might be an important improvement to existing transplantation protocols to other retinal neurons, as well.

### Statistical analysis

All graphs were generated by GraphPad PRISM 9 (GraphPad software, San Diego, CA). Statistical significance was determined using an unpaired Welch’s *t*-test. All data are presented as average ± standard deviation (SD). Where normally distributed, the data were analyzed with a *p* value of <0.05 (**\****p* < 0.05, **\*\****p* < 0.01, **\*\*\****p* < 0.001, **\*\*\*\****p* < 0.0001) as a statistically significant.

## Acknowledgments

This work was supported by NEI/NIH U24 (PB), Gilbert Family Foundation (PB), VRP FTTSA VR220053 (PB), Gilbert Family Foundation (PB). The authors would like to thank Dr. Don Zack for providing the Brn3b-tdTomato hESCs.

## Author contributions

MHP, VVM, MGK and PB initiated the project and designed experiments. VVM, MHP, JA performed confocal microscopy, immunohistochemistry experiments. JDR differentiated donor hRGCs. JA analyzed the images of microglia cells. EK assembled a single-cell atlas for mouse retina. CA performed scanning laser ophthalmoscopy. VVM and PB performed cell transplantation, wrote the manuscript draft. PB supervised the project. All authors reviewed and contributed to editing the manuscript.

## Data availability statement

The original code generated in this study is available on GitHub (https://github.com/mcrewcow/Malechka_et_al_2025), and the atlases constructed and utilized in the manuscript are deposited to CELLxGENE (https://cellxgene.cziscience.com/collections/7dd7e8a1-f7c2-4213-9f23-dd52afb7ee16).

## Supplementary figure legends

**Figure 1.** Elimination of donor human retinal ganglion cells (hRGCs) by primary microglia *in vitro* on day 0 (**A**) and day 1 (**B**).

**Supplementary Table 1.**

| **GSE** | **DOI** | **Condition** |
| --- | --- | --- |
| GSE123758 | Not published_3, as in CELLxGENE | Mouse healthy adult retina atlas |
| GSE125708 | 10.1073/pnas.1821122116<br>(https://pubmed.ncbi.nlm.nih.gov/30988181/) | Mouse healthy adult retina atlas |
| GSE132229 | 10.1073/pnas.1915571116<br>(https://pubmed.ncbi.nlm.nih.gov/31843893/) | Mouse healthy adult retina atlas |
| GSE137398 | 10.1016/j.neuron.2019.11.006<br>(https://pubmed.ncbi.nlm.nih.gov/31784286/) | Mouse healthy adult retina atlas |
| GSE137828 | 10.1016/j.neuron.2019.11.006<br>(https://pubmed.ncbi.nlm.nih.gov/31784286/) | Mouse healthy adult retina atlas |
| GSE153674 | PMID: 33088174<br>(https://pubmed.ncbi.nlm.nih.gov/33088174/) | Mouse healthy adult retina atlas |
| GSE164044 | 10.1016/i.neuron.2019.08.002<br>(https://pubmed.ncbi.nlm.nih.gov/31493975/) | Mouse healthy adult retina atlas |
| GSE169262 | 10.1038/s41467-021-27924-y<br>(https://pubmed.ncbi.nlm.nih.gov/35017532/) | Mouse healthy adult retina atlas |
| GSE176204 | 10.1016/i.celrep.2023.112889<br>(https://pubmed.ncbi.nlm.nih.gov/37527036/) | Mouse healthy adult retina atlas |
| GSE183572 | 10.1038/s42003-022-03676-3<br>(https://www.nature.com/articles/s42003-022-03676-3) | Mouse healthy adult retina atlas |
| GSE199317 | 10.1038/s41590-023-01437-w<br>(https://pubmed.ncbi.nlm.nih.gov/36807640/) | Mouse healthy adult retina atlas |
| GSE204880 | 10.1038/s42003-023-05300-4<br>(https://pubmed.ncbi.nlm.nih.gov/37670124/) | Mouse healthy adult retina atlas |
| GSE205123 | 10.1016/j.ygeno.2023.110644<br>(https://pubmed.ncbi.nlm.nih.gov/37279838/) | Mouse healthy adult retina atlas |
| GSE220626 | 10.3389/fcell.2023.1252547<br>(https://pubmed.ncbi.nlm.nih.gov/37691820/) | Mouse healthy adult retina atlas |
| GSE232470 | 10.1186/s40478-023-01580-3<br>(https://pubmed.ncbi.nlm.nih.gov/37226256/) | Mouse healthy adult retina atlas |
| GSE236389 | Not published_1, as in CELLxGENE | Mouse healthy adult retina atlas |
| GSE239347 | 10.1073/pnas.2314698120<br>(https://pubmed.ncbi.nlm.nih.gov/38064509/) | Mouse healthy adult retina atlas |
| GSE243100 | 10.2337/db23-0368<br>(https://pubmed.ncbi.nlm.nih.gov/37976214/) | Mouse healthy adult retina atlas |
| GSE243413 | 10.1016/i.isci.2024.109916<br>(https://pubmed.ncbi.nlm.nih.gov/38812536/) | Mouse healthy adult retina atlas |
| GSE251717 | 10.1101/2024.03.18.585452<br>(https://pubmed.ncbi.nlm.nih.gov/38562864/) | Mouse healthy adult retina atlas |
| GSE263877 | 10.1038/s41467-024-50033-5<br>(https://pubmed.ncbi.nlm.nih.gov/39009597/) | Mouse healthy adult retina atlas |
| GSE137398 | 10.1016/i.neuron.2019.11.006<br>(https://pubmed.ncbi.nlm.nih.gov/31784286/) | Mouse optic nerve crush retina |
| GSE199317 | 10.1038/s41590-023-01437-w<br>(https://pubmed.ncbi.nlm.nih.gov/36807640/) | Mouse optic nerve crush retina |
| GSE201254 | 10.1016/i.neuron.2022.06.002<br>(https://pubmed.ncbi.nlm.nih.gov/35767994/) | Mouse optic nerve crush retina |
| GSE206622 | 10.1016/j.neuron.2022.06.022<br>(https://pubmed.ncbi.nlm.nih.gov/35952672/) | Mouse optic nerve crush retina |
| GSE206623 | 10.1016/j.neuron.2022.06.022<br>(https://pubmed.ncbi.nlm.nih.gov/35952672/) | Mouse optic nerve crush retina |
| GSE206624 | 10.1016/j.neuron.2022.06.022<br>(https://pubmed.ncbi.nlm.nih.gov/35952672/) | Mouse optic nerve crush retina |
| GSE206625 | 10.1016/j.neuron.2022.06.022<br>(https://pubmed.ncbi.nlm.nih.gov/35952672/) | Mouse optic nerve crush retina |
| GSE223530 | 10.1073/pnas.2305947121<br>(https://pubmed.ncbi.nlm.nih.gov/38289952/) | Mouse optic nerve crush retina |
| GSE228627 | 10.1101/2023.11.17.567614<br>(https://pubmed.ncbi.nlm.nih.gov/38014168/) | Mouse optic nerve crush retina |
| GSE232470 | 10.1186/s40478-023-01580-3<br>(https://pubmed.ncbi.nlm.nih.gov/37226256/) | Mouse optic nerve crush retina |
| GSE241268 | 10.1038/s41467-024-46428-z<br>(https://pubmed.ncbi.nlm.nih.gov/38467611/) | Mouse optic nerve crush retina |
| GSE248537 | 10.1038/s41467-024-46428-z<br>(https://pubmed.ncbi.nlm.nih.gov/38467611/) | Mouse optic nerve crush retina |
| GSE226456 | 10.1016/i.celrep.2023.112889<br>(https://pubmed.ncbi.nlm.nih.gov/37527036/) | Mouse MB-induced glaucoma retina |
| GSE247776 | 10.1038/s41467-024-50050-4<br>(https://pubmed.ncbi.nlm.nih.gov/39080269/) | Mouse MB-induced glaucoma retina |
| not published | not published, as in CELLxGENE (Dong Feng Chen Lab) | Mouse MB-induced glaucoma retina |
| GSE118614 | 10.1016/i.neuron.2019.04.010<br>(https://pubmed.ncbi.nlm.nih.gov/31128945/) | Mouse healthy developing retina |
| GSE122466 | 10.1242/dev.178103<br>(https://pubmed.ncbi.nlm.nih.gov/31399471/) | Mouse healthy developing retina |
| GSE139904 | 10.1126/sciadv.abj9846<br>(https://pubmed.ncbi.nlm.nih.gov/34757798/) | Mouse healthy developing retina |
| GSE186069 | 10.1038/s41586-024-07069-w<br>(https://pubmed.ncbi.nlm.nih.gov/38947520/) | Mouse healthy developing retina |
| GSE172101 | 10.1242/dev.199399<br>(https://pubmed.ncbi.nlm.nih.gov/34143204/) | Mouse healthy developing retina |
| GSE202734 | 10.1016/j.celrep.2023.112237<br>(https://pubmed.ncbi.nlm.nih.gov/36924502/) | Mouse healthy developing retina |
| GSE222956 | 10.1016/j.celrep.2023.112237<br>(https://pubmed.ncbi.nlm.nih.gov/36924502/) | Mouse healthy developing retina |
| GSE228590 | 10.1038/s41586-024-07069-w<br>(https://pubmed.ncbi.nlm.nih.gov/38947520/) | Mouse healthy developing retina |
| GSE245542 | 10.1016/i.celrep.2024.114291<br>(https://pubmed.ncbi.nlm.nih.gov/38823017/) | Mouse healthy developing retina |
| GSE252861 | 10.1016/i.isci.2024.110100<br>(https://pubmed.ncbi.nlm.nih.gov/38947520/) | Mouse healthy developing retina |

## References

Quigley, H.A., and Broman, A.T. (2006). The number of people with glaucoma worldwide in 2010 and 2020. Br. J. Ophthalmol. 90, 262. 10.1136/bjo.2005.081224.

Cook, C., and Foster, P. (2012). Epidemiology of glaucoma: what’s new? Can. J. Ophthalmol. J. Can. d’Ophtalmol. 47, 223–226. 10.1016/j.jcjo.2012.02.003.

Newman, N.J., and Biousse, V. (2004). Hereditary optic neuropathies. Eye 18, 1144–1160. 10.1038/sj.eye.6701591.

Patil, A.D., Biousse, V., and Newman, N.J. (2022). Ischemic Optic Neuropathies: Current Concepts. Ann. Indian Acad. Neurol. 25, S54–S58. 10.4103/aian.aian_533_22.

Klistorner, A., Sriram, P., Vootakuru, N., Wang, C., Barnett, M.H., Garrick, R., Parratt, J., Levin, N., Raz, N., Walt, A.V. der, et al. (2014). Axonal loss of retinal neurons in multiple sclerosis associated with optic radiation lesions. Neurology 82, 2165–2172. 10.1212/wnl.0000000000000522.

Zhang, K.Y., and Johnson, T.V. (2021). The internal limiting membrane: Roles in retinal development and implications for emerging ocular therapies. Exp Eye Res 206, 108545. 10.1016/j.exer.2021.108545.

Baranov, P., and Oswald, J. (2018). Retinal Ganglion Cell Replacement: A Bridge to the Brain. In Regenerative Medicine and Stem Cell Therapy for the Eye., pp. 193–206. 10.1007/978-3-319-98080-5_8.

Shen, J., Xiao, R., Bair, J., Wang, F., Vandenberghe, L.H., Dartt, D., Baranov, P., and Ng, Y.S.E. (2018). Novel engineered, membrane-localized variants of vascular endothelial growth factor (VEGF) protect retinal ganglion cells: a proof-of-concept study. Cell death & disease 9, 1371. 10.1038/s41419-018-1049-0.

Soucy, J.R., Aguzzi, E.A., Cho, J., Gilhooley, M.J., Keuthan, C., Luo, Z., Monavarfeshani, A., Saleem, M.A., Wang, X.-W., Wohlschlegel, J., et al. (2023). Retinal ganglion cell repopulation for vision restoration in optic neuropathy: a roadmap from the RReSTORe Consortium. Mol. Neurodegener. 18, 64. 10.1186/s13024-023-00655-y.

Singh, M.S., Park, S.S., Albini, T.A., Canto-Soler, M.V., Klassen, H., MacLaren, R.E., Takahashi, M., Nagiel, A., Schwartz, S.D., and Bharti, K. (2020). Retinal stem cell transplantation: Balancing safety and potential. Prog. Retin. Eye Res. 75, 100779. 10.1016/j.preteyeres.2019.100779.

Zhang, J., Wu, S., Jin, Z.-B., and Wang, N. (2021). Stem Cell-Based Regeneration and Restoration for Retinal Ganglion Cell: Recent Advancements and Current Challenges. Biomolecules 11, 987. 10.3390/biom11070987.

Venugopalan, P., Wang, Y., Nguyen, T., Huang, A., Muller, K.J., and Goldberg, J.L. (2016). Transplanted neurons integrate into adult retinas and respond to light. Nat. Commun. 7, 10472. 10.1038/ncomms10472.

Hertz, J., Qu, B., Hu, Y., Patel, R.D., Valenzuela, D.A., and Goldberg, J.L. (2013). Survival and Integration of Developing and Progenitor-Derived Retinal Ganglion Cells following Transplantation. Cell Transplant. 23, 855–872. 10.3727/096368913x667024.

Vrathasha, V., Nikonov, S., Bell, B.A., He, J., Bungatavula, Y., Uyhazi, K.E., and Chavali, V.R.M. (2022). Transplanted human induced pluripotent stem cells-derived retinal ganglion cells embed within mouse retinas and are electrophysiologically functional. iScience 25, 105308. 10.1016/j.isci.2022.105308.

Rashid, K., Akhtar-Schaefer, I., and Langmann, T. (2019). Microglia in Retinal Degeneration. Front. Immunol. 10, 1975. 10.3389/fimmu.2019.01975.

Karlstetter, M., Scholz, R., Rutar, M., Wong, W.T., Provis, J.M., and Langmann, T. (2015). Retinal microglia: Just bystander or target for therapy? Prog. Retin. Eye Res. 45, 30–57. 10.1016/j.preteyeres.2014.11.004.

Chen, M., and Xu, H. (2015). Parainflammation, chronic inflammation, and age-related macular degeneration. J. Leucoc. Biol. 98, 713–725. 10.1189/jlb.3ri0615-239r.

Pan, L., Cho, K.-S., Wei, X., Xu, F., Lennikov, A., Hu, G., Tang, J., Guo, S., Chen, J., Kriukov, E., et al. (2023). IGFBPL1 is a master driver of microglia homeostasis and resolution of neuroinflammation in glaucoma and brain tauopathy. Cell Rep. 42, 112889. 10.1016/j.celrep.2023.112889.

Vermes, I., Haanen, C., Steffens-Nakken, H., and Reutellingsperger, C. (1995). A novel assay for apoptosis Flow cytometric detection of phosphatidylserine expression on early apoptotic cells using fluorescein labelled Annexin V. J. Immunol. Methods 184, 39–51. 10.1016/0022-1759(95)00072-i.

Kägi, D., Vignaux, F., Ledermann, B., Bürki, K., Depraetere, V., Nagata, S., Hengartner, H., and Golstein, P. (1994). Fas and Perforin Pathways as Major Mechanisms of T Cell-Mediated Cytotoxicity. Science 265, 528–530. 10.1126/science.7518614.

Suda, T., Hashimoto, H., Tanaka, M., Ochi, T., and Nagata, S. (1997). Membrane Fas Ligand Kills Human Peripheral Blood T Lymphocytes, and Soluble Fas Ligand Blocks the Killing. J. Exp. Med. 186, 2045–2050. 10.1084/jem.186.12.2045.

Hohlbaum, A.M., Moe, S., and Marshak-Rothstein, A. (2000). Opposing Effects of Transmembrane and Soluble FAS Ligand Expression on Inflammation and Tumor Cell Survival. J. Exp. Med. 191, 1209–1220. 10.1084/jem.191.7.1209.

Gregory, M.S., Hackett, C.G., Abernathy, E.F., Lee, K.S., Saff, R.R., Hohlbaum, A.M., Moody, K.L., Hobson, M.W., Jones, A., Kolovou, P., et al. (2011). Opposing Roles for Membrane Bound and Soluble Fas Ligand in Glaucoma-Associated Retinal Ganglion Cell Death. PLoS ONE 6, e17659. 10.1371/journal.pone.0017659.

Rogge, M., Yin, X.-T., Godfrey, L., Lakireddy, P., Potter, C.A., Rosso, C.R.D., and Stuart, P.M. (2015). Therapeutic Use of Soluble Fas Ligand Ameliorates Acute and Recurrent Herpetic Stromal Keratitis in Mice. Investig. Opthalmology Vis. Sci. 56, 6377. 10.1167/iovs.15-16588.

Li, N.-L., Nie, H., Yu, Q.-W., Zhang, J.-Y., Ma, A.-L., Shen, B.-H., Wang, L., Bai, J., Chen, X.-H., Zhou, T., et al. (2004). Role of soluble Fas ligand in autoimmune diseases. World J. Gastroenterol. 10, 3151–3156. 10.3748/wjg.v10.i21.3151.

Fligor, C.M., Huang, K.-C., Lavekar, S.S., VanderWall, K.B., and Meyer, J.S. (2020). Differentiation of retinal organoids from human pluripotent stem cells. Methods Cell Biol. 159, 279–302. 10.1016/bs.mcb.2020.02.005.

Oswald, J., Kegeles, E., Minelli, T., Volchkov, P., and Baranov, P. (2021). Transplantation of miPSC/mESC-derived retinal ganglion cells into healthy and glaucomatous retinas. Mol. Ther. - Methods Clin. Dev. 21, 180–198. 10.1016/j.omtm.2021.03.004.

Soucy, J.R., Todd, L., Kriukov, E., Phay, M., Malechka, V.V., Rivera, J.D., Reh, T.A., and Baranov, P. (2023). Controlling donor and newborn neuron migration and maturation in the eye through microenvironment engineering. Proc. Natl. Acad. Sci. 120, e2302089120. 10.1073/pnas.2302089120.

Young, K., and Morrison, H. (2018). Quantifying Microglia Morphology from Photomicrographs of Immunohistochemistry Prepared Tissue Using ImageJ. J. Vis. Exp. : JoVE, 57648. 10.3791/57648.

Alt, C., and Lin, C.P. (2012). In vivo quantification of microglia dynamics with a scanning laser ophthalmoscope in a mouse model of focal laser injury. Ophthalmic Technol. XXII, 820907-820907–820909. 10.1117/12.909141.

O’Koren, E.G., Yu, C., Klingeborn, M., Wong, A.Y.W., Prigge, C.L., Mathew, R., Kalnitsky, J., Msallam, R.A., Silvin, A., Kay, J.N., et al. (2019). Microglial Function Is Distinct in Different Anatomical Locations during Retinal Homeostasis and Degeneration. Immunity 50, 723–737.e7. 10.1016/j.immuni.2019.02.007.

Carelli, V., Morgia, C.L., Ross-Cisneros, F.N., and Sadun, A.A. (2017). Optic neuropathies: the tip of the neurodegeneration iceberg. Hum. Mol. Genet. 26, R139–R150. 10.1093/hmg/ddx273.

Johnson, T.V., Bull, N.D., Hunt, D.P., Marina, N., Tomarev, S.I., and Martin, K.R. (2010). Neuroprotective Effects of Intravitreal Mesenchymal Stem Cell Transplantation in Experimental Glaucoma. Investig. Opthalmology Vis. Sci. 51, 2051. 10.1167/iovs.09-4509.

Chamling, X., Sluch, V.M., and Zack, D.J. (2016). The Potential of Human Stem Cells for the Study and Treatment of Glaucoma. Investig. Ophthalmol. Vis. Sci. 57, ORSFi1–ORSFi6. 10.1167/iovs.15-18590.

Silva-Junior, A.J. da, Mesentier-Louro, L.A., Nascimento-dos-Santos, G., Teixeira-Pinheiro, L.C., Vasques, J.F., Chimeli-Ormonde, L., Bodart-Santos, V., Carvalho, L.R.P. de, Santiago, M.F., and Mendez-Otero, R. (2021). Human mesenchymal stem cell therapy promotes retinal ganglion cell survival and target reconnection after optic nerve crush in adult rats. Stem Cell Res. Ther. 12, 69. 10.1186/s13287-020-02130-7.

Chang, E.E., and Goldberg, J.L. (2011). Glaucoma 2.0: neuroprotection, neuroregeneration, neuroenhancement. Ophthalmology 119, 979–986. 10.1016/j.ophtha.2011.11.003.

Wareham, L.K., Risner, M.L., and Calkins, D.J. (2020). Protect, Repair, and Regenerate: Towards Restoring Vision in Glaucoma. Curr. Ophthalmol. Rep. 8, 301–310. 10.1007/s40135-020-00259-5.

Murenu, E., Gerhardt, M.-J., Biel, M., and Michalakis, S. (2022). More than meets the eye: The role of microglia in healthy and diseased retina. Front. Immunol. 13, 1006897. 10.3389/fimmu.2022.1006897.

Fan, W., Huang, W., Chen, J., Li, N., Mao, L., and Hou, S. (2022). Retinal microglia: Functions and diseases. Immunology 166, 268–286. 10.1111/imm.13479.

Fadok, V.A., Savill, J.S., Haslett, C., Bratton, D.L., Doherty, D.E., Campbell, P.A., and Henson, P.M. (1992). Different populations of macrophages use either the vitronectin receptor or the phosphatidylserine receptor to recognize and remove apoptotic cells. J. Immunol. 149, 4029– 4035. 10.4049/jimmunol.149.12.4029.

Martin, S.J., Reutelingsperger, C.P., McGahon, A.J., Rader, J.A., Schie, R.C. van, LaFace, D.M., and Green, D.R. (1995). Early redistribution of plasma membrane phosphatidylserine is a general feature of apoptosis regardless of the initiating stimulus: inhibition by overexpression of Bcl-2 and Abl. J. Exp. Med. 182, 1545–1556. 10.1084/jem.182.5.1545.

Logue, S.E., Elgendy, M., and Martin, S.J. (2009). Expression, purification and use of recombinant annexin V for the detection of apoptotic cells. Nat. Protoc. 4, 1383–1395. 10.1038/nprot.2009.143.

Gerke, V. (2001). Annexins And Membrane Organisation In The Endocytic Pathway. Cell. Mol. Biol. Lett. 6, 204.

Gerke, V., and Moss, S.E. (2002). Annexins: From Structure to Function. Physiol. Rev. 82, 331–371. 10.1152/physrev.00030.2001.

Koopman, G., Reutelingsperger, C., Kuijten, G., Keehnen, R., Pals, S., and Oers, M. van (1994). Annexin V for flow cytometric detection of phosphatidylserine expression on B cells undergoing apoptosis. Blood 84, 1415–1420. 10.1182/blood.v84.5.1415.bloodjournal8451415.

Krishnan, A., Fei, F., Jones, A., Busto, P., Marshak-Rothstein, A., Ksander, B.R., and Gregory-Ksander, M. (2016). Overexpression of Soluble Fas Ligand following Adeno-Associated Virus Gene Therapy Prevents Retinal Ganglion Cell Death in Chronic and Acute Murine Models of Glaucoma. J. Immunol. 197, 4626–4638. 10.4049/jimmunol.1601488.

Tanaka, M., Itai, T., Adachi, M., and Nagata, S. (1998). Downregulation of Fas ligand by shedding. Nat. Med. 4, 31–36. 10.1038/nm0198-031.

Gregory, M.S., Repp, A.C., Holhbaum, A.M., Saff, R.R., Marshak-Rothstein, A., and Ksander, B.R. (2002). Membrane Fas Ligand Activates Innate Immunity and Terminates Ocular Immune Privilege. J. Immunol. 169, 2727–2735. 10.4049/jimmunol.169.5.2727.

Matsumoto, H., Murakami, Y., Kataoka, K., Notomi, S., Mantopoulos, D., Trichonas, G., Miller, J.W., Gregory, M.S., Ksander, B.R., Marshak-Rothstein, A., et al. (2015). Membrane-bound and soluble Fas ligands have opposite functions in photoreceptor cell death following separation from the retinal pigment epithelium. Cell Death Dis. 6, e1986. 10.1038/cddis.2015.334.

Gregory-Ksander, M., Perez, V.L., Marshak-Rothstein, A., and Ksander, B.R. (2019). Soluble Fas ligand blocks destructive corneal inflammation in mouse models of corneal epithelial debridement and LPS induced keratitis. Exp. Eye Res. 179, 47–54. 10.1016/j.exer.2018.10.013.

Krishnan, A., Kocab, A.J., Zacks, D.N., Marshak-Rothstein, A., and Gregory-Ksander, M. (2019). A small peptide antagonist of the Fas receptor inhibits neuroinflammation and prevents axon degeneration and retinal ganglion cell death in an inducible mouse model of glaucoma. J. Neuroinflammation 16, 184. 10.1186/s12974-019-1576-3.

Mendes-Jorge, L., Ramos, D., Luppo, M., Llombart, C., Alexandre-Pires, G., Nacher, V., Melgarejo, V., Correia, M., Navarro, M., Carretero, A., et al. (2009). Scavenger function of resident autofluorescent perivascular macrophages and their contribution to the maintenance of the blood-retinal barrier. Investig. Ophthalmol. Vis. Sci. 50, 5997–6005. 10.1167/iovs.09-3515.

Sasaki, Y., Ohsawa, K., Kanazawa, H., Kohsaka, S., and Imai, Y. (2001). Iba1 Is an Actin-Cross-Linking Protein in Macrophages/Microglia. Biochem. Biophys. Res. Commun. 286, 292–297. 10.1006/bbrc.2001.5388.

