## Supplementary figures and images for "Improvement of human donor retinal ganglion cell survival through modulation of microglia"

### Supplemental Figure 1

A B

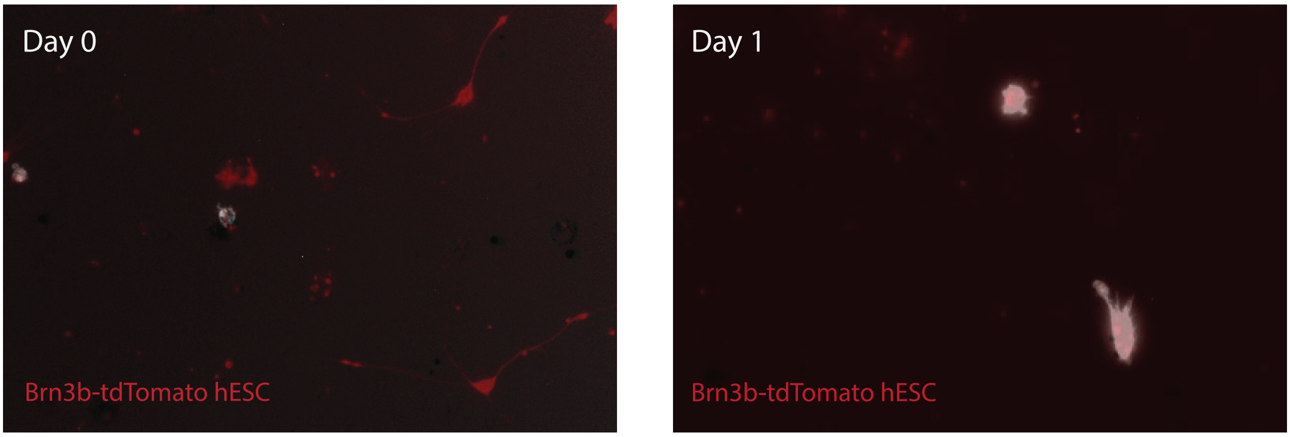


Supplementary Figure 1. Malechka VV et al. 2025
